# KLP-7/Kinesin-13 orchestrates axon-dendrite checkpoints for polarized trafficking in neurons

**DOI:** 10.1101/2023.08.24.554630

**Authors:** Swagata Dey, Nitish Kumar, Jessica Feldman, Anindya Ghosh-Roy

**Affiliations:** Cellular and Molecular Neuroscience, National Brain Research Centre, Manesar, Gurugram, Haryana, India; Present affiliation: Huck Institutes of Life Sciences, Pennsylvania State University, PA, USA; School of Humanities and Sciences, Stanford University, CA, USA

**Keywords:** Kinesin13, microtubule depolymerization, PVD neuron, *C. elegans*, neuronal polarity

## Abstract

Neurons are compartmentalized cells with spatiotemporal distinction of anatomy and molecular repertoire. Microtubule organization in the neuron is crucial for its polarized structure and composition. Microtubule dynamics are differentially optimized in the axons and dendrites by the interplay between the microtubule-stabilizing and destabilizing factors. It is unclear how the destabilizing factors are important for developing and maintaining neuronal polarity.

We investigated the function of KLP-7, a microtubule depolymerizing motor from the Kinesin-13 family, in the compartmentalization of axons and dendrites using the PVD neurons in *Caenorhabditis elegans*. In the absence of KLP-7, axonal proteins such as RAB-3 and SAD-1 were mislocalized to dendrites, suggesting a disruption in axon-dendrite compartmentalization. Notably, this phenomenon was independent of other depolymerizing factors like EFA-6, highlighting the specific role of KLP-7 in this process. We observed a reduced rate of microtubule polymerization and an altered polarity of microtubules in the PVD major dendrite due to the loss of *klp-7*. Additionally, the deletion of *klp-7* led to the formation of ectopic neurites from the cell body and the ectopic localization of UNC-44/Ankyrin-G, a protein associated with the axon initial segment (AIS), to the dendrites. Additionally, live imaging of GFP::KLP-7 revealed that KLP-7 is more dynamic in the dendrites as compared to the axon. These observations indicate that the precise dynamics of KLP-7 in neurites are crucial for maintaining distinct microtubule polymerization in the axons and dendrites, thereby influencing neuronal polarity.

Our findings shed light on the pivotal role of KLP-7/Kinesin-13 in the establishment of axon-dendrite checkpoints, which in turn impact the polarized trafficking of cellular components within neurons.

## Introduction

The polarized structure and composition of the neurons enable directional information transfer in a neural circuit. Microtubules are one of the crucial determiners of neuronal polarization (Barnes and Polleux, 2009; Marín *et al*., 2010; Kuijpers and Hoogenraad, 2011; Yogev and Shen, 2017). Dynamics and organization of microtubules define neuronal migration, selective intracellular trafficking, organelle positioning, signaling hubs, and force generators in the developing neurons (Tolić-Nørrelykke, 2008; Hirokawa *et al*., 2010; De Forges *et al*., 2012; Subramanian and Kapoor, 2012). Microtubules are made of α and β tubulin heterodimers equilibrating between phases of polymerization and depolymerization that define their polarity, dynamics, and organization during neuronal morphogenesis (Sakakibara *et al*., 2013; Horio *et al*., 2014; Kapitein and Hoogenraad, 2015; Kelliher *et al*., 2019; Gudimchuk and McIntosh, 2021).

In the context of neuronal polarity, regulators of microtubule stabilization have been well investigated (Baas *et al*., 1991, 2016; Witte *et al*., 2008; Polleux and Snider, 2010) for example, MAP2C and TRIM46 stabilize the microtubule array and are important for the axon initiation (Dehmelt *et al*., 2006; Van Beuningen *et al*., 2015). Similarly, microtubule assembly regulator Collapsin Response Mediator Protein 2 (CRMP2) regulates the compartmentalization of axons and dendrites (Arimura *et al*., 2004; Yoshimura *et al*., 2005; Maniar *et al*., 2012). Microtubule destabilizers like Stathmin/Op18 are locally inactivated in the nascent axon during development (Watabe-Uchida *et al*., 2006). Similarly, a microtubule severing protein, Katanin is finely regulated for the proper axon development (Karabay *et al*., 2004; Yu *et al*., 2005; Qiang *et al*., 2006). Conditional loss of microtubule depolymerizing KIF2/Kinesin-13 in the rodent hippocampus and KLP-7 in *Caenorhabditis elegans* caused the formation of ectopic branches and loss of axon-dendrite compartmentalization (Homma *et al*., 2003, 2018; Chen *et al*., 2011; Ghosh-Roy *et al*., 2012; Puri *et al*., 2021). The roles and mechanisms of microtubule depolymerizers in the process of neuronal polarization are yet to be investigated in detail.

PVD neurons are highly polarized neurons with a well-defined axon and highly stereotyped dendritic arbor. These are mechanosensory neurons for proprioception, harsh touch, and cold temperature sensation (Way and Chalfie, 1989; Chatzigeorgiou *et al*., 2010; Albeg *et al*., 2011). Previous studies have elucidated the role of various regulators including Ankyrin (UNC-44), CRMP2 (UNC-33), UNC-119, Patronin (PTRN-1), Ninein (NOCA-2), and TIAM-1 in the process of neuronal polarity (Maniar *et al*., 2012; He *et al*., 2020, 2022; Lin *et al*., 2022). Loss of function of these genes caused aberrant microtubule organization and mislocalization of the axonal cargoes into the dendrites (Maniar *et al*., 2012; He *et al*., 2020, 2022; Lin *et al*., 2022). As these molecules participate during various stages of the PVD development and regulate microtubule organization differently, it is difficult to ascertain how microtubule depolymerization participates in axon-dendrite compartmentalization.

In this study, we have found specific role of KLP-7/Kinesin-13 in proper compartmentalization of axons and dendrites in the PVD neurons. In the absence of KLP-7, the axonal cargoes such as RAB-3 and SAD-1 populate the dendrites disrupting the axon-dendrite compartmentalization. Using reporter of microtubule dynamics, EBP-2::GFP, we have found that KLP-7 maintains the microtubule polarity and organization in the major dendrite of the PVD neuron. Also, KLP-7 is important for the specific accumulation of the AIS component, UNC-44 in the AIS as an ectopic accumulation of UNC-44 occurs in the dendrites of *klp-7(0)* mutant. Our study suggests that more dynamics of KLP-7 in the dendrites than axon prevents ubiquitous or ectopic enrichment of AIS components by regulating the microtubule organization in the dendrites of the PVD neurons.

## Results

### KLP-7/Kinesin-13 regulates the axon-dendrite compartmentalization in the PVD neurons

KLP-7 is a motor of the Kinesin-13 family required for the dynamic instability of the microtubules (Desai *et al*., 1999; Srayko *et al*., 2005). Conditional knockout of KIF2 (mammalian ortholog of Kinesin-13) in the hippocampi resulted in the formation of abnormal neurites in the dentate granule cells which were not differentiated into the axons and dendrites in terms of their molecular constitution (Ogawa and Hirokawa, 2015; Homma *et al*., 2018). Similar anatomical defects were also observed in the touch neurons of *C. elegans*, where the loss of the *klp-7* gene resulted in the formation of ectopic neurites with axonal markers (Puri *et al*., 2021). The mechanism by which Kinesin-13 regulates the axonal and dendritic identities is not clear.

We investigated aspects of neuronal polarity in the loss of function mutant of *klp-7* (Kinesin-13). We checked the distribution of the axonal marker, RAB-3 as a reporter for neuronal polarity in the PVD neuron (Figure 1A). PVD neurons of *C. elegans* have a well-defined axon running ventrally and fasciculated with the ventral nerve cord and a highly stereotyped dendritic arbor with orthogonal branches (Albeg *et al*., 2011; Sundararajan *et al*., 2019). The mCherry::RAB-3 reporter appears as punctae mostly in the ventral nerve cord region in the control (wildtype) background (Figure 1A). Unlike the wildtype, mCherry::RAB-3 was also present in the dendrites of the *klp-7* null mutant (Figure 1B-C). Estimation of the dendritic branches containing mCherry::RAB-3 showed an accumulation in 11.5% of the dendritic branches of *klp-7(0)* as compared to 3.5% in the wildtype (Figure 1D). Though the loss of *klp-7* caused localization of mCherry::RAB-3 throughout the PVD neuron including the minor dendrite, it was more pronounced in the distal region of the major dendrite where *klp-7* null mutant had 29.4% dendrites with mCherry::RAB-3 as compared to 11.8% of the wildtype (Figure S1A). This accumulation was prominent in the tertiary branches of *klp-7(0)* with 33.2% occupancy (Figure S1B). Interestingly, quaternary branches of the *klp-7* null mutant also showed mislocalization of mCherry::RAB-3 (Figure S1B). Moreover, the precedence of mCherry::RAB-3 in the major dendrite as observed by the particle density was also increased in the null mutant of *klp-7* (Figure 1E). It could have resulted due to an increased anterograde transport in the presence of stable microtubules in the mutant.

**Figure 1:**
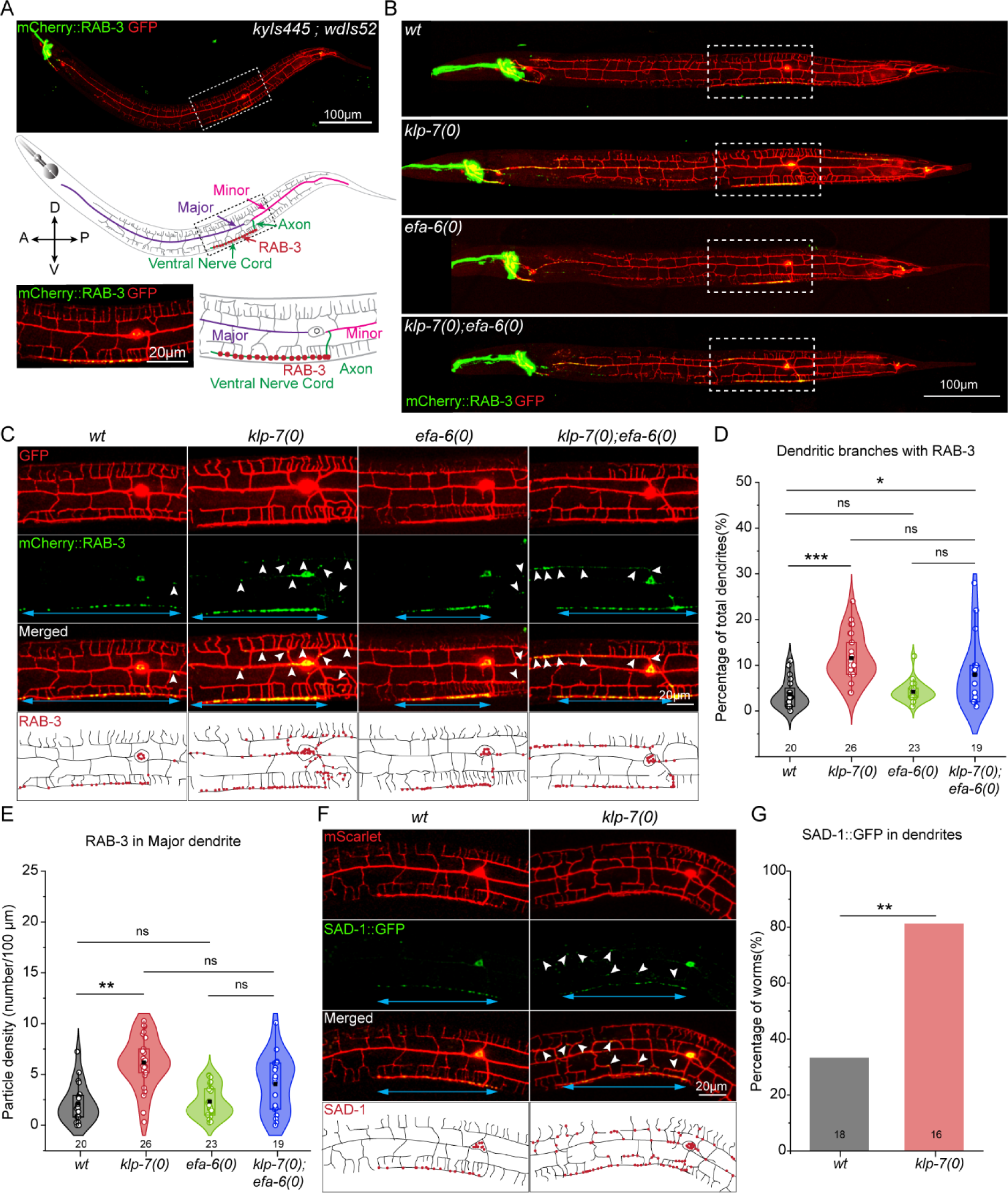
KLP-7 is required for the proper axon-dendrite compartmentalization in the PVD neurons. A. Representative images and schematics of the PVD neuron expressing soluble GFP and mCherry::RAB-3, with the region around the cell body (dashed box) magnified below. Schematic depicts crosshair reference points of body symmetry (A-anterior, P-posterior, D-dorsal, V-ventral). B-C. Representative images of the whole PVD neuron expressing soluble GFP and mCherry::RAB-3 in the wildtype (wt), and loss of function mutants of *klp-7(0)*, *efa-6(0)*, and double mutant of *klp-7(0)* and *efa-6(0)* with the regions around the cell body (dashed box) magnified and schematized in (C). The axonal and dendritic distributions of mCherry::RAB-3 in (C) are marked by the blue arrows and white arrowheads, respectively. Schematic in (C) shows the distribution of mCherry::RAB-3 (red dots) in the dendritic arbor of various mutants. D. Quantification of the dendritic branches showing accumulation of mCherry::RAB-3 in the wildtype (wt), and loss of function mutants of *klp-7(0)*, *efa-6(0)*, and double mutant of *klp-7(0)* and *efa-6(0)* normalized to the total number of dendritic branches represented as a percentage. Comparison of means was done using ANOVA and Bonferroni Test. p < 0.05*, 0.001***, ns (not significant). Number of animals assessed (n) is mentioned along the X-axis. E. Number of mCherry::RAB-3 particles in the major dendrite normalized to the length of the dendrite in the wildtype (wt), and loss of function mutants of *klp-7(0)*, *efa-6(0)*, and double mutant of *klp-7(0)* and *efa-6(0)*. Comparison of means was done using ANOVA and Bonferroni Test. p < 0.01**, ns (not significant). Number of animals assessed (n) is mentioned along the X-axis. F-G. Representative images and schematic (F), and quantification (G) of SAD-1::GFP in the dendrites of the wildtype (wt), and loss of function mutants of *klp-7(0)*. The axonal and dendritic distribution of SAD-1::GFP are marked by blue arrows and white arrowheads, respectively. Schematic in (F) shows the distribution of SAD-1::GFP (red dots) in the dendritic arbor of wildtype (wt) and *klp-7(0)*. Proportion of the animals showing dendritic accumulation of SAD-1::GFP with respect to the total animals assessed is represented as a percentage. Comparison of datasets was done using Fisher’s exact test. p <0.01**. Number of animals assessed (n) is mentioned along the X-axis.

A similar phenotype was observed for another synaptic molecule SAD-1 which also gets mislocalized to the dendrites in the absence of *klp-7* (Figure 1F). Around 80% of the animals showed this phenotype (Figure 1G). This indicated that KLP-7 limits the entry of axonal cargo by compartmentalization as well as anterograde traffic into the dendrites. Though mCherry::RAB-3 particle density in the minor dendrite and the axon showed a tendency to increase due to the loss of *klp-7*, it was not significantly perturbed (Figure S1C-D). This indicated that KLP-7 has a differential requirement in the polarized transport in PVD compartments.

As KLP-7 is a microtubule catastrophe factor, we asked if any other microtubule catastrophe factor would also regulate the axon-dendrite compartmentalization. We investigated another mutant that has a loss of function of *efa-6*. EFA6 is a cortical protein that is an exchange factor for ARF6 GTPase required in membrane trafficking and actin regulation (Franco *et al*., 1999). The *C. elegans* ortholog, EFA-6 has a microtubule depolymerizing domain that causes the catastrophe of the microtubules near the cell cortex (O’Rourke *et al*., 2010). Loss of *EFA-6* causes the cells to have long microtubules and has been phenotypically correlated to spindle defects, embryonic lethality, and *dlk-1-*independent axon regeneration (O’Rourke *et al*., 2010; Chen *et al*., 2015; Qu *et al*., 2019). Unlike the loss of function mutant of *klp-7*, null mutant of *efa-6* did not mislocalize RAB-3 to the dendrites (Figure 1B-C). Measurement of the number of dendrites with RAB-3 and distribution of RAB-3 in the primary dendrite of the PVD neuron of *efa-6(0)* were equivalent to those of wildtype (Figure 1D-E, S1A-B). The axon-dendrite distinction seems to be preserved in this mutant like the wildtype. Furthermore, we did not observe a synergism between *klp-7* and *efa-6* in the double mutant as compared to the *klp-7* mutant alone (Figure 1B-E, S1A-B). Only in the posterior dendrite (minor), the effect of *klp-7(0)* appeared to be suppressed by *efa-6(0)* which further strengthened the hypothesis of the differential requirement of KLP-7 in PVD neurites (Figure S1A). This also indicated that the role of KLP-7 in neuronal polarity is highly specific even though both KLP-7 and EFA-6 are microtubule destabilizing factors.

### Microtubule organization by KLP-7 is a determiner of the axon-dendrite compartmentalization

Microtubule organization in the neurons is important for the differential trafficking of the cargoes in the axonal vs dendritic compartments (Kapitein *et al*., 2010; Kapitein and Hoogenraad, 2011; Maeder *et al*., 2014). Previous studies have established the causal effect of microtubule stabilization on axon specification by pharmacological perturbation of the microtubules through Paclitaxel to induce multiple neurites with axonal identity (Witte *et al*., 2008). Paclitaxel decreases depolymerization, promotes assembly, and increases the stabilization of the microtubules (Xiao *et al*., 2006). On the other hand, microtubule destabilizers like Nocodazole and Colchicine could reduce the formation of dendrite like processes (Witte *et al*., 2008; Puri *et al*., 2021).

To understand if KLP-7 mediated microtubule organization defines the dendritic identity, we used pharmacological disruption of microtubules by Colchicine. Previously, Colchicine exposure has been used to suppress the effects of microtubule-stabilizing mutations in *C. elegans* (Kirszenblat *et al*., 2013; Puri *et al*., 2021). We used a concentration of 1mM colchicine which has been shown to affect the microtubule dynamics and function of the gentle touch neurons (Chalfie and Thomson, 1982; Bounoutas *et al*., 2011; Puri *et al*., 2021). On exposure to 1mM Colchicine, we observed no difference in the dendritic distribution of RAB-3 in the wildtype, however, in the *klp-7* loss of function mutant, the prevalence of RAB-3 in the dendrites decreased significantly with the exposure of the Colchicine (Figure 2A-B). This indicates that KLP-7 maintains an optimal levels of microtubule dynamics in the dendrites to prevent the entry of axonal cargoes like RAB-3 into the dendrites. Though the reduction in neuronal polarity defect in *klp-7(0)* by Colchicine was barely perceptible in branches, it was mostly evident in the medial and proximal areas of the PVD neuron marginally, in the tertiary dendrites (Figure S2A-B). This was a partial change as a significant number of *klp-7(0)* dendrites still accumulated RAB-3 in the presence of Colchicine as compared to its treated wildtype control (Figure 2B-D). Due to Colchicine treatment, density of mCherry::RAB-3 particles decreased in the wildtype axons whereas it was unaffected in the *klp-7(0)* mutant (Figure 2E). This indicated that KLP-7 dependent organization of microtubules is one of the factors determining the axon-dendrite compartmentalization and polarized transport. KLP-7 may have additional role that determines the axon-dendrite compartmentalization.

**Figure 2:**
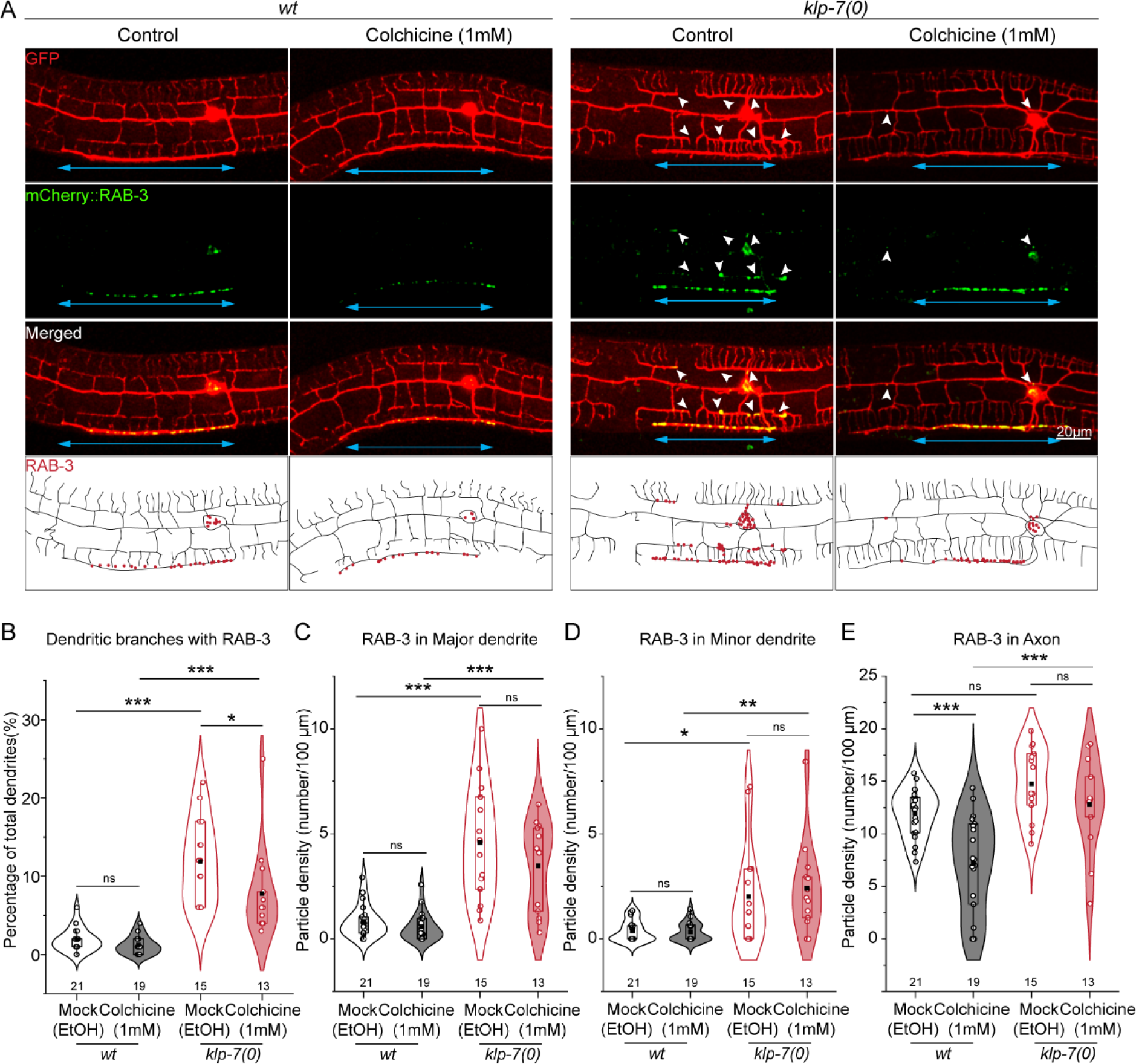
KLP-7-dependent microtubule organization defines the axon-dendrite compartmentalization. A. Representative images of the wildtype (wt) and *klp-7(0)* mutant following the treatment with Mock (Ethanol) and Colchicine (1 mM). The axonal and dendritic distributions of mCherry::RAB-3 are marked by the blue arrows and white arrowheads, respectively. Schematic shows the distribution of mCherry::RAB-3 (red dots) in the dendritic arbor of wildtype (wt) and *klp-7(0)* upon Colchicine treatment. B. Percentage of the dendritic branches with respect to the total number of dendritic branches showing accumulation of mCherry::RAB-3 in the wildtype (wt), and *klp-7(0)*mutant in mock and Colchicine treated conditions. Comparison of means was done using ANOVA and Bonferroni Test. p < 0.05*, 0.01**, 0.001***, ns (not significant). Number of animals assessed (n) is mentioned along the X-axis. C-E. Quantification of particle density of mCherry::RAB-3 in the major (C), minor (D), and axon (E) in the wildtype (wt) and *klp-7(0)* mutant upon Colchicine treatment with their respective mock controls. Comparison of means was done using ANOVA and Bonferroni Test and p > 0.05. Number of animals assessed (n) is mentioned along the X-axis.

### Kinesin-13 regulates the relative orientation and distribution of dynamic microtubules in the major dendrite

The two major aspects of microtubule organization that define the neuronal polarity or polarized trafficking in the neuron are microtubule dynamics and orientation. Kinesin-13 is a key determinant of dynamic instability (Desai *et al*., 1999; Ogawa *et al*., 2017; Trofimova *et al*., 2018). Puri et.al. 2021 showed that KLP-7 maintains a mixed orientation of the microtubules by making them more dynamic in the posterior process of the PLM neurons (Puri *et al*., 2021). This process in the PVD dendrite is additionally regulated by UNC-116, PTRN-1, NOCA-2, TIAM-1, and UNC-119 (He *et al*., 2020, 2021; Liang *et al*., 2020; Lin *et al*., 2022). The role of KLP-7 in maintaining the dendritic microtubule organization is yet to be characterized.

To understand the microtubule organization, we observed the dynamics of the plus tips of the microtubules using the EBP-2::GFP marker. EBP-2::GFP binds to the plus ends of microtubules and shows comet-like movement in a time lapse acquisition (Stepanova *et al*., 2003; Ghosh-Roy *et al*., 2012). To assess the EBP-2::GFP dynamics, we generated the kymographs in the PVD neurites with origin at the cell body (Figure 3A). In the wildtype neurons, the major dendrite showed minus end-out microtubules with the comets moving towards the cell body whereas the minor dendrite and the axon showed plus end-out orientation of the microtubules (Figure 3B). This is similar to previous studies measuring the EBP-2::GFP comets in PVD neurons (Maniar *et al*., 2012; Harterink *et al*., 2018). In the *klp-7(0)* mutant, the major dendrite showed a significant number of comets that are moving in the opposite direction as also visible in the kymographs (Figure 3B). We estimated the percentage of EBP-2::GFP comets in either direction for each kymograph. Based on the percentage of the EBP-2::GFP comets the major dendrite of the wildtype had a minus end-out orientation of the microtubules whereas the mutant has a mixed orientation (Figure 3C). On the other hand, *klp-7* loss of function did not perturb the orientation of the microtubules in the minor dendrite or the axon (Figure 3C, S3A-B). Both the minor dendrite and the axon of the wildtype neurons have plus end-out polarity of the microtubules that was unperturbed in the mutant (Figure 3C, S3A-B). This suggested that KLP-7 is specifically required for the microtubule polarity, in the anterior dendrite (major).

**Figure 3:**
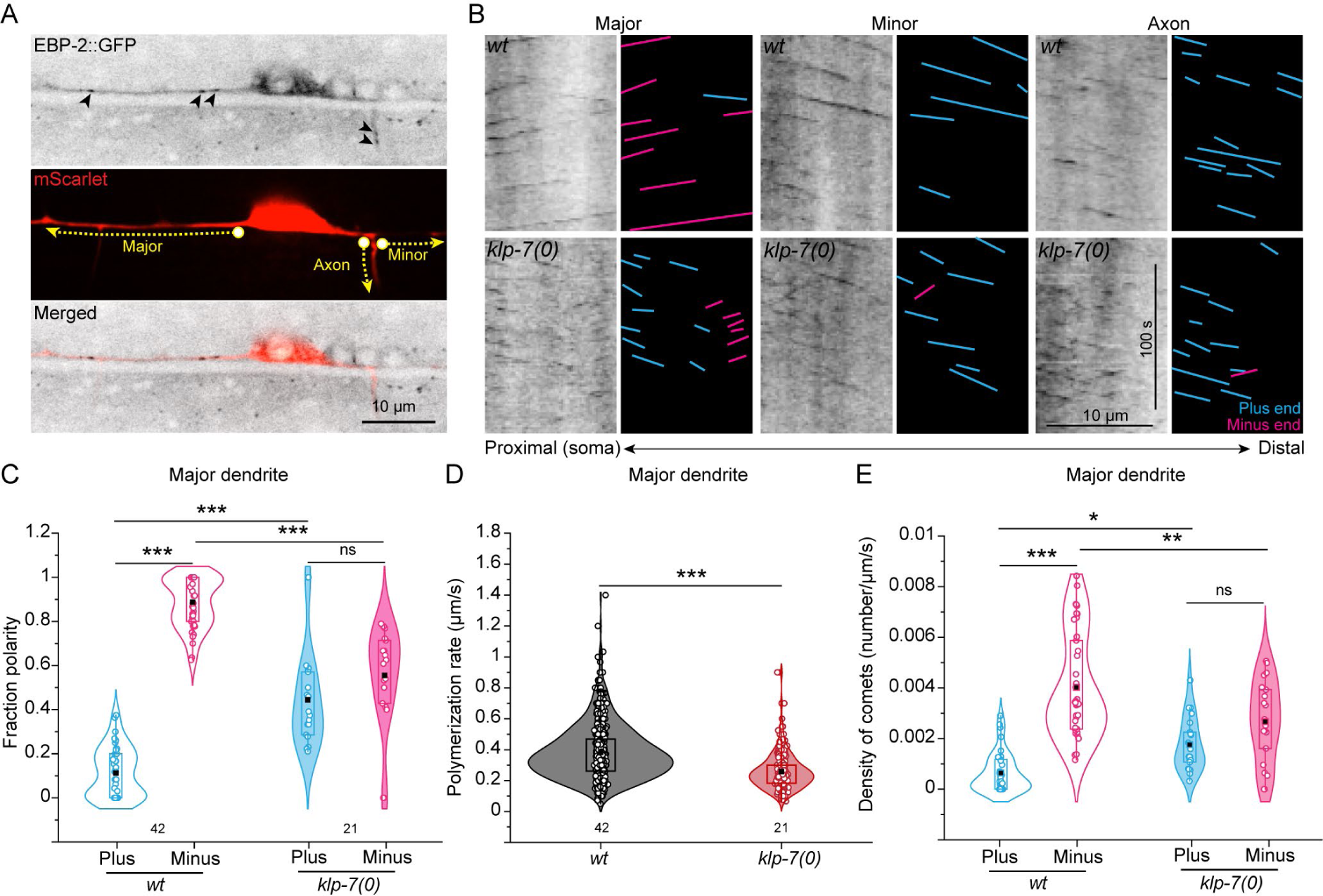
Microtubule polarity and dynamics in the PVD dendrites perturbed by loss of *klp-7* function. A. Representative images of endogenously expressed EBP-2::GFP and PVD neuron expressing mScarlet. Different neurites used for assessing EBP-2::GFP comet dynamics are depicted with yellow segmented lines. The origin of the segmented line is marked by a circle which refers to the proximal region in the kymographs. B. Kymographs with their respective schematics showing the trajectories of EBP-2::GFP comet in the PVD major dendrite, minor dendrite, and the axonal compartments of wildtype (wt) and *klp-7(0)*. Schematics show the orientation of the comets in the plus-end-out (cyan) and minus-end-out (magenta) directions. C. Relative orientation of the microtubules is represented as a fraction polarity of EBP-2::GFP comets in the plus-end-out (cyan) and minus-end-out directions (magenta) in the PVD major dendrite of wildtype (wt) and *klp-7(0)*. Comparison of means was done using ANOVA and Bonferroni Test. p < 0.001***. Number of animals assessed (n) is mentioned along the X-axis. D. Rates of polymerization were quantified as the covered distance divided by the duration of the EBP-2::GFP comets in the PVD major dendrite of wildtype (wt) and *klp-7(0)*. Comparison of means was done using ANOVA and Bonferroni Test. p < 0.001***. Number of animals assessed (n) is mentioned along the X-axis. E. Densities of EBP-2::GFP comets were quantified for plus and minus end-out microtubules in the PVD major dendrite of wildtype (wt) and *klp-7(0)*. Comparison of means was done using ANOVA and Bonferroni Test. p < 0.05*, 0.01**, 0.001***. Number of animals assessed (n) is mentioned along the X-axis.

Unlike wildtype dendrites, robust polymerization events were diminished which was evident in the rates of microtubule polymerization which was reduced in the mutant (Figure 3D, S3D-E). All the neurites including the major dendrite, minor dendrite, and axon showed a decreased rate of polymerization in the absence of *klp-7* (Figure 3D, S3D-E). This could have resulted from the increased microtubule pauses in the absence of KLP-7-dependent depolymerization as observed for *Drosophila* microtubules due to accumulation of CLASP or loss of Mini Spindles (Msps) from their plus ends (Li *et al*., 2011; Moriwaki and Goshima, 2016). Interestingly, we observed distribution of the EBP-2::GFP comets was altered in the *klp-7* loss of function mutant (Figure S3E). In the wildtype, EBP-2::GFP comets are present in the primary dendrites and the axon whereas in the *klp-7(0)* mutant these comets also went into secondary branches and sometimes to tertiary branches (Figure S3E). This could be due to the ingression of stabilized microtubules in the higher-order branches. Consequentially, it may increase the anterograde traffic of the RAB-3 vesicles to the higher-order dendrites.

We also quantified the density of comets in either direction which is the number of comets per unit length of the major dendrite and per unit time of acquisition. Instead of an overall decrease in the number of comets due to microtubule stabilization, the density of minus end-out comets was decreased, and plus end-out comets was increased in the *klp-7(0)* mutant in its primary major dendrite (Figure 3E). Also, the plus end out microtubules in *klp-7(0)* major dendrite seemed to be closer to the cell body (Figure 3B). These results indicate that Kinesin-13/KLP-7 is limiting the entry of plus end-out microtubules in the major dendrite of PVD neurons. Previous results have elucidated that increased stabilization of microtubules in the *klp-7(0)* results in the loss of the minus end proteins like PTRN-1 (Puri *et al*., 2021). We checked the presence of the minus end protein PTRN-1::tagRFP in the PVD dendrites to understand if the minus ends are perturbed due to the loss of the *klp-7* gene. The density of the PTRN-1::tagRFP was comparable between the wildtype and *klp-7(0)* neurons (Figure S3F-G). Based on comet density and PTRN-1::tagRFP punctae, we concluded that KLP-7 is maintaining the microtubule polarity possibly by forming a dendritic checkpoint that regulates the entry of the plus end out microtubules into the major dendrite.

### Axon Initial Segment (AIS) protein UNC-44 accumulates in the dendrites of *klp-7(0)*

Previous studies have explored the dynamic microtubules in the PVD dendrites and deciphered AnkyrinG (UNC-44), CRMP2 (UNC-33), UNC-119, Kinesin-1 (UNC-116), Patronin (PTRN-1), and Ninein (NOCA-2) as the major regulators of microtubule dynamics (Maniar *et al*., 2012; Yan *et al*., 2013; Taylor *et al*., 2015; He *et al*., 2020, 2022). Some of the factors like UNC-44, UNC-33, and UNC-119 (Figure 4A) also regulate the polarized trafficking into the dendrites (He *et al*., 2020). It is unclear how KLP-7 is participating in defining the axon-dendrite compartmentalization and microtubule orientation in the dendrites.

**Figure 4:**
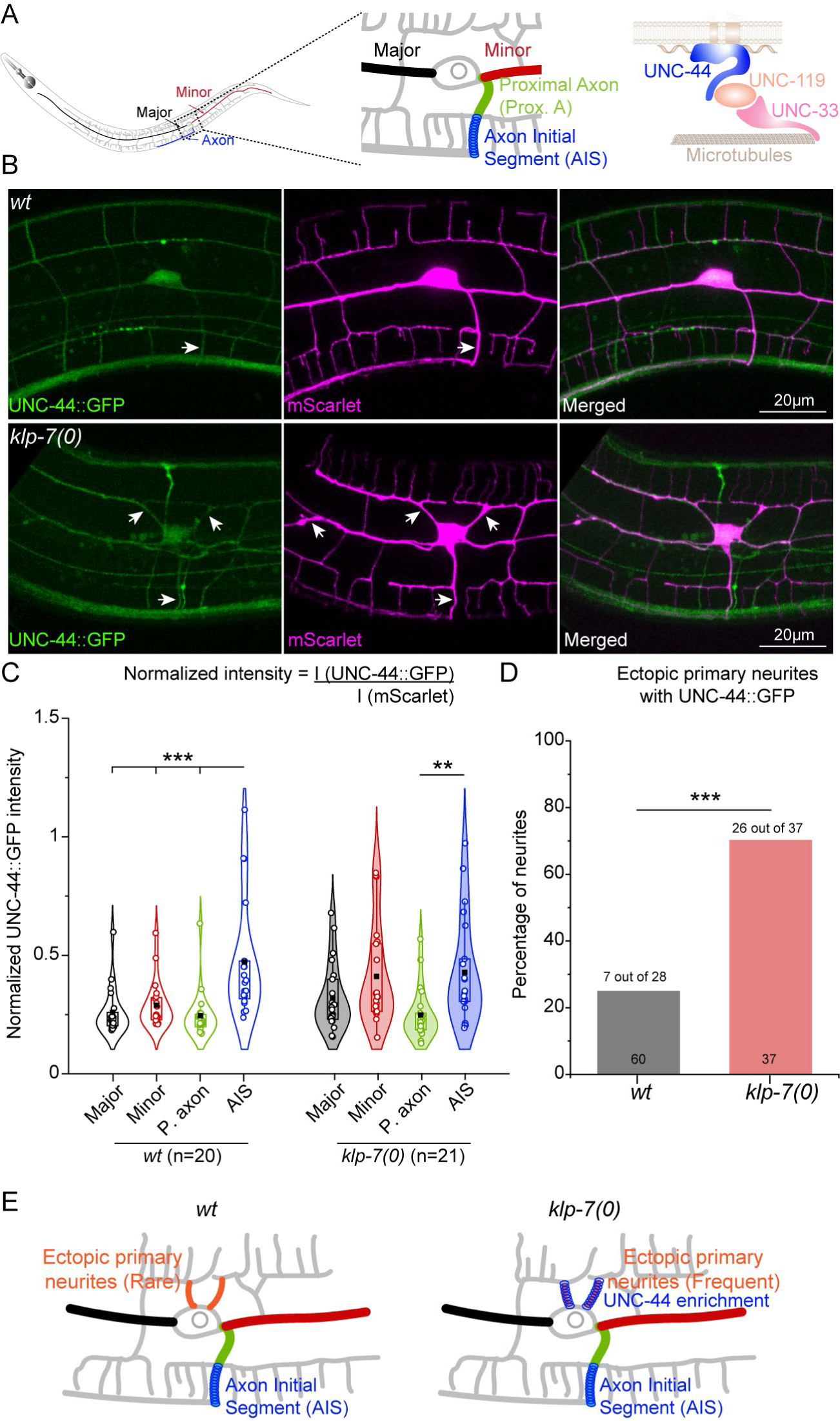
Specific enrichment of Axon Initial Segment protein, UNC-44 is dependent on *klp-7* function. A. Schematic showing the major, minor, proximal axon, and axon initial segment compartments to assess the distribution of UNC-44::GFP in a typical PVD neuron magnified in the middle panel. Right panel shows the constitution of axon initial segment adapted from He *et al*., 2020. B. Representative images of the PVD neuron in wildtype and *klp-7(0)* expressing mScarlet with single copy expression of UNC-44::GFP under its endogenous promoter. White arrows depict the enrichment of UNC-44::GFP in various neurites of the PVD neuron. C. Normalized intensities of UNC-44::GFP to a constitutive PVD reporter mScarlet in the neurites of PVD neurons of wildtype and *klp-7(0)*. Comparison of means was done using ANOVA and Bonferroni Test. p<0.01**, 0.001***. Number of animals assessed (n) is mentioned along the X-axis. D. Ectopic branches emanating from the cell bodies of wildtype and *klp-7(0)* neurons were assessed for their enrichment and quantified as the percentage of neurites with UNC-44::GFP with respect to total neurites assessed. Number of assessed PVD neurons are mentioned adjacent to the X-axis. Proportion of the ectopic neurites showing enrichment of UNC-44::GFP is mentioned on top of the bar plot. Statistical comparison was done using Fisher’s test p<0.001***. E. Schematic of PVD neurons with ectopic branches in wildtype and *klp-7(0)* showing prominent enrichment of UNC-44 in these ectopic branches of the *klp-7(0)* mutant.

To explore the role of KLP-7 in the maintenance of the AIS, we observed single CRISPR mediated GFP knockin at the N terminus of UNC-44, in the PVD neurons (He *et al*., 2020). Based on the molecular constitution and function of the mammalian AIS, a homologous structure has been defined in the laterodistal region of the PVD axon (Figure 4A). UNC-44::GFP expressed under its own promoter has ubiquitous expression and is present in all the neurites of PVD neurons (Figure 4B). Normalized and mean intensity quantification of UNC-44::GFP in the major, minor, proximal axon (P. axon), and AIS showed a remarkable enrichment in the AIS region (Figure 4B-C, S4A). The intensity was lowest in the proximal axon region (Figure 4B-C, S4A). In the *klp-7(0)* mutant, the bonafide AIS also showed a similar degree of enrichment (Figure S4A). As protein densities could be different as per the cytosolic constitution in dendrites and AIS, we normalized the intensity of UNC-44::GFP to a constitutive marker mScarlet in the PVD (Figure 4C). Relative intensities of UNC-44::GFP to mScarlet also showed a similar trend of enrichment of UNC-44::GFP at the AIS as compared to the other regions of the wildtype neurons (Figure 4C). However, in the loss of *klp-7* function mutant, both major and minor dendrites of the PVD neuron showed comparable levels to their respective AIS (Figure 4C). This indicated that differential enrichment of UNC-44 in the AIS as compared to the other neurites requires *klp-7*. Moreover, the expression levels of UNC-44::GFP in the cell bodies of the PVD neuron in the wildtype and *klp-7* mutant strengthens this observation (Figure S4B).

We further investigated how intensities of UNC-44::GFP in the *klp-7(0)* dendrites were relatively higher. Interestingly, loss of *klp-7* also results in the formation of ectopic branches that are emanating from the cell body (Figure 4B, arrowheads). The occurrence of such ectopic branches was quite lower in the wildtype. As compared to the ectopic branches in the wildtype neurons, the *klp-7(0)* neurons showed an enrichment comparable to that of the AIS (Figure 4B, D). Around 70% of the ectopic branches of the *klp-7(0)* showed AIS-like enrichment as compared to 25% of the wildtype ectopic branches (Figure 4D-E). The formation of AIS coincides with the formation of a polarized array of microtubules by increased stabilization and fasciculation which are codependent to ensure polarized sorting of the cargoes between axonal and dendritic compartments (Fréal *et al*., 2019). Increased microtubule stabilization in *klp-7* null mutant may enrich UNC-44 in a non-specific manner to PVD neurites which causes aberrant sorting of RAB-3, and plus end out microtubules.

### Differential regulation of microtubule organization by Kinesin-13 in PVD compartments

Neuronal polarization is initiated by the axon differentiation which depends on microtubule stabilization (Witte *et al*., 2008). In *Drosophila* neurons, perturbation in the microtubule polarity could direct the formation of ectopic diffusion barriers (Thyagarajan *et al*., 2022). To correlate the formation of AIS and the function of KLP-7, we investigated the localization and dynamics of GFP::KLP-7b isoform in the wildtype neurons. This isoform was shown to rescue the anatomical defects in the touch receptor neurons (Puri *et al*., 2021).

We investigated the presence and mobility of GFP::KLP-7b in the major dendrite, minor dendrite, proximal axon (P. axon), and the AIS of the PVD neuron (Figure 5A). Localization of GFP::KLP-7 was equivalent in all these compartments (Figure 5A, lower panel). Normalization with the constitutive marker mScarlet further validated this observation (Figure 5B). All the assessed compartments showed an equivalent quantity of GFP::KLP-7b (Figure 5B). Mobility of KLP-7 has been correlated to its activity in the gentle touch neurons (Puri *et al*., 2021). We further assessed the dynamics of GFP::KLP-7b in these compartments (Figure 5C) by Fluorescence Recovery After Photobleaching (FRAP) assay. The FRAP assay revealed that the mobility of GFP::KLP-7b is significantly lesser in the axon initial segment with respect to the other compartments (Figure 5C-D). Quantification of the maximum recovery from the recovery profiles showed a considerably small mobile fraction in the AIS as compared to the major dendrite, minor dendrite, and p.axon (Figure S5A). Moreover, a comparison of the halftimes of the recovery profiles also showed a delayed recovery in the AIS as compared to the major dendrite (Figure S5B). This is in line with the previous observations in wildtype PLM neurons where activity or mobility of KLP-7 was comparatively lower in the anterior process akin to the axon (Puri *et al*., 2021). The differential activity of KLP-7 in different compartments of the neuron allows these compartments to acquire a distinct microtubule organization (Puri *et al*., 2021).

**Figure 5:**
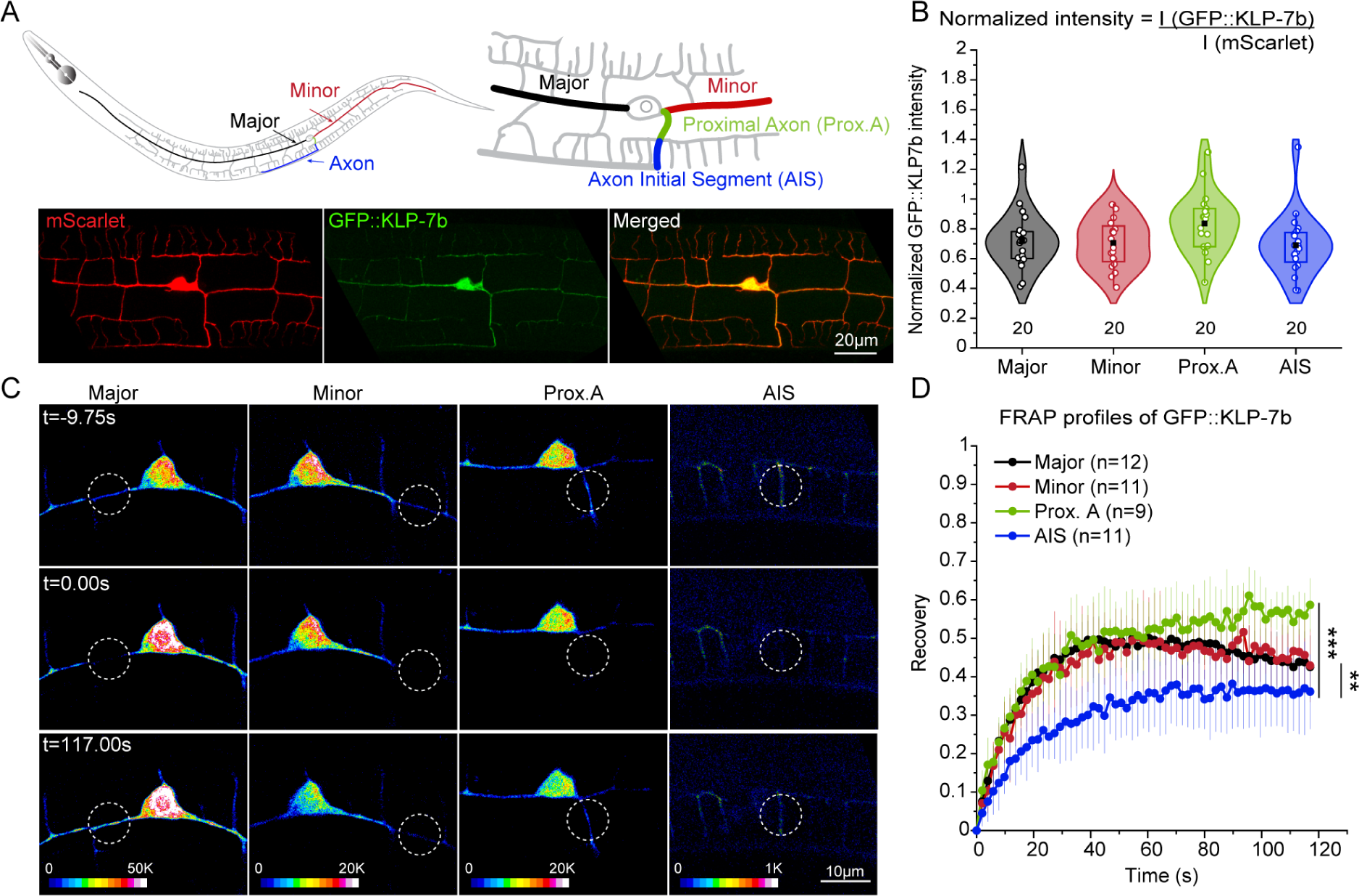
Differential activity of KLP-7 in the PVD axonal and dendritic compartments. A. Schematic and representative image showing the major, minor, proximal axon, and axon initial segment compartments to assess the distribution of GFP::KLP-7b isoform (5ng extrachromosomal array) in the PVD neuron. B. Intensities of GFP::KLP-7b in the various compartments of the PVD neuron normalized to the constitutive marker mScarlet (10ng extrachromosomal array). The datasets were not found to be significantly different. Number of animals assessed (n) is mentioned along the X-axis. C-D. Representative images (C) of Fluorescence Recovery After Photobleaching assay of GFP::KLP-7b in various compartments quantified as recovery profiles (D). Number of animals assessed (n) is mentioned in the legend. Comparison of maximum recovery was done using ANOVA and Bonferroni Test. p < 0.01**, 0.001***. Number of animals assessed (n) is mentioned in the legend.

**Figure 6.**
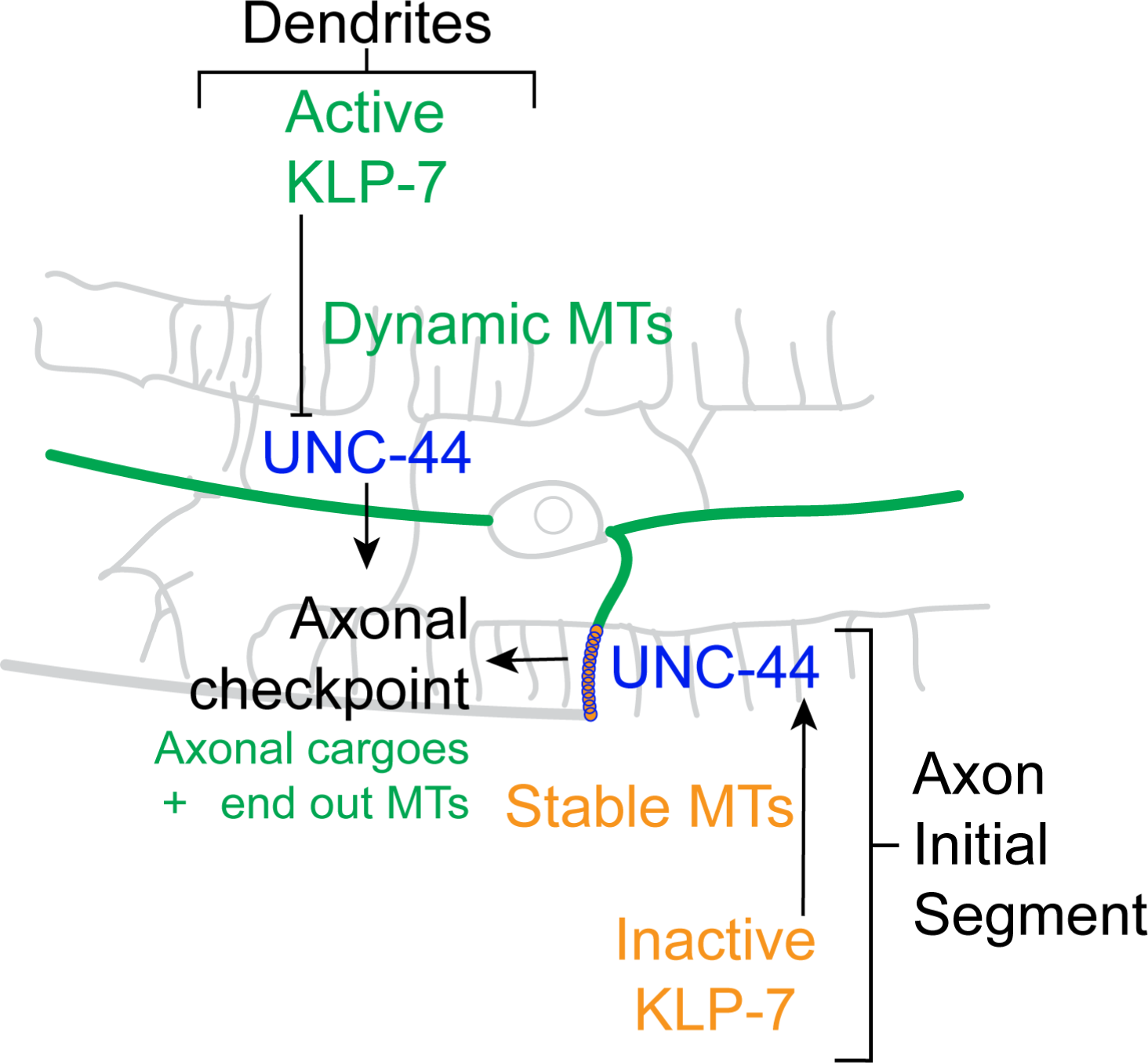
KLP-7 mediated axonal checkpoint determining axonal sorting of cargoes and microtubules. Differential activity of KLP-7 in the axon and dendrites of PVD neurons maintain stable and dynamic microtubules (MTs), respectively in these compartments. Loss of *klp-7* correlated to ectopic enrichment of Ankyrin (UNC-44) in the dendrites also resulted in stable microtubules, increased dendritic presence of axonal cargoes and plus end out microtubules in the dendrites.

To validate this observation, we compared the rates of polymerization obtained from EBP-2::GFP comets (Figure 2) in the dendrites and axon. Rates of polymerization were higher in both the major and minor dendrite as compared to the axon in the wildtype neurons (Figure S5C). This suggests that microtubules are more dynamic in the dendrites as compared to the axon. In the *klp-7(0)* mutant, not only the rates of polymerization were decreased globally, but also the distinction between the axon and dendrites became obsolete (Figure S5C). This suggests that relative microtubule dynamics in the various neurites are correlated to the differential activity of KLP-7 in these compartments.

## Discussion

Our study demonstrates that KLP-7/Kinesin-13 plays a crucial role in regulating the compartmentalization of axons and dendrites. We have identified a dual role for KLP-7, which appears to converge in maintaining neuronal polarity in PVD neurons. The first function involves the maintenance of microtubule polarity and distribution in the major dendrite, while the second function involves restricting the enrichment of axon initial segment components in the dendrites. Our findings suggest that KLP-7 exhibits higher activity in the dendrites compared to the axons. This higher activity leads to increased microtubule depolymerization in the dendrites, which in turn prevents the accumulation of UNC-44, an axon initial segment protein in the dendrites. The accumulation of UNC-44 might further direct plus end out microtubule organization in the axon.

### KLP-7/Kinesin-13 dependent microtubule organization regulates the axon dendrite compartmentalization

Neuronal polarization commences with axon specification (Dotti *et al*., 1988; Higgs and Das, 2022) determined by microtubule stabilization and bundling (Witte *et al*., 2008; Van Beuningen *et al*., 2015). Stability of the microtubules imparts strength to the growing axon, provides platform for other cytoskeletal interactions, promotes selective trafficking of molecules, and generate force for changes in morphology during growth (Hawkins *et al*., 2010; Kapitein and Hoogenraad, 2011). It is unclear how microtubule catastrophe factors which also contribute to dynamic instability of microtubules participate in the neuronal polarization.

Loss of Kinesin-13/KIF2/KLP-7 causes an aberrant arborization of neurons with defects in axon-dendrite compartmentalization (Homma *et al*., 2018; Puri *et al*., 2021). Conditional KIF2 knockout mice also showed behavioral defects like epilepsy (Homma *et al*., 2018). This study also showed similar defects in the PVD neurons but the underlying mechanism was elusive. Microtubule stabilization in the loss of function of KLP-7 (Kinesin-13) could lead to formation of multiple axon like neurites during development, similar to that observed upon Taxol treatment (Witte *et al*., 2008). We showed other microtubule depolymerizers like EFA-6 did not affect the axon-dendrite compartmentalization. However, we could partially suppress the defect of *klp-7* null mutant by Colchicine treatment indicating additional roles of KLP-7 besides microtubule stabilization in preserving the axon-dendrite compartmentalization.

KLP-7 is a microtubule depolymerizing enzyme and under the Wnt signaling its differential activity in the PLM neurites regulates its microtubule organization (Desai *et al*., 1999; Puri *et al*., 2021). but it was unclear how KLP-7 organizes the microtubules in the PVD neurons. We showed that loss of *klp-7* resulted in decreased polymerization and a mixed orientation of microtubules in the major dendrite which typically has minus end out microtubule polarity. In other neurites with plus end out polarity i.e. minor and axon, the polymerization dynamics were affected but the polarity of the microtubules was unchanged by *klp-7* deletion. In the *klp-7* null mutant, apart from increase in the plus end out EBP-2::GFP comets, we observed a concomitant decrease in the minus end out microtubules which rules out the possibility of specific action of KLP-7 on plus end out microtubules in the major dendrite. It could be an effect of global disruption of intraneuronal trafficking of cargoes including axonal cargoes and microtubules.

### Differential activity of KLP-7 regulates formation and integrity of the diffusion barriers in the PVD neurons

Previous studies have explored neuronal polarity in the PVD neurons and have found role of UNC-44, UNC-33, UNC-119, TIAM-1, PTRN-1 and NOCA-2 (Maniar *et al*., 2012; He *et al*., 2020, 2022; Lin *et al*., 2022). In loss of function mutants of these genes, besides the mislocalization of axonal cargoes in the dendrites, microtubule polarity in the neurites was also perturbed (Maniar *et al*., 2012; He *et al*., 2020, 2022; Lin *et al*., 2022). Additional mechanisms like mislocalization of UNC-104 to dendrites, defects in Kinesin-1 mediated microtubule sliding, loss of cytoskeletal interactions, and possible defects in microtubule nucleation were also associated to these mutants (Maniar *et al*., 2012; He *et al*., 2020, 2022; Lin *et al*., 2022). It was unclear how KLP-7 coordinates with these molecules to regulate the microtubule organization and neuronal polarity.

As per our hypothesis, change in the microtubule polarity in the major dendrite in the *klp-7(0)* could be due to aberrant intracellular traffic, we investigated the axon initial segment protein, UNC-44. Although, the protein was ubiquitously localized to all neurites of PVD, normalized intensity of this protein in the wildtype PVD neurons showed a marked enrichment in the bonafide Axon Initial Segment defined as per the molecular constitution akin to mammalian AIS (Maniar *et al*., 2012; He *et al*., 2020). Loss of *klp-7* affected the selective enrichment in the AIS instead we observed an increase in relative intensity of UNC-44::GFP in the dendrites of PVD neurons. This was suggestive of ectopic accumulation of the diffusion barrier proteins in the dendrites. This was promiscuous in the ectopic branches emanating from the cell bodies of PVD neurons. Though this anatomical feature was rare in wildtype animals, these branches showed excess of UNC-44::GFP in the deletion mutant of *klp-7*. This result corroborated previous observation of mosaic arbors in *Drosophila* class IV ddaC neurons upon Patrnonin RNAi which also showed the formation of ectopic diffusion barriers (Thyagarajan *et al*., 2022). An increased precedence of plus end out microtubules near the cell body in *klp-7(0)* PVD dendrites exalted their tendency to accumulate the diffusion barrier proteins. Increased dynamics of GFP::KLP-7b in the dendrites than in the axon supports this idea. KLP-7 activity in the dendrites forms check points in the dendrites to regulate the microtubule polymerization, polarity, and distribution into higher order dendrites. This further regulates the formation and integrity of diffusion barrier thereby affecting the selective intracellular transport. Overall, our study sheds light on the intricate mechanisms underlying the regulation of axon-dendrite compartmentalization through KLP-7 mediated microtubule regulation and AIS formation and integrity.

## Materials and Methods

### *C. elegans* culturing, constructs, and transgenesis

The worms were cultured on the Nematode Growth Medium (NGM) seeded with OP50 bacteria at 20⁰C. Mutants and transgenics were crossed and homozygosity of mutant and transgene alleles was confirmed by genotyping and phenotyping. The mutant and transgenes are mentioned in Table S1. The allele *kyIs445 [pdes-2::mCherry::RAB-3 (0.5 ng/µL); pdes-2::SAD-1::GFP (2 ng/µL); podr-1::DsRed (30 ng/µL)]* is a gift from Cory Bargmann lab, and *ebp-2*(*wow47*[*ebp-2::gfp::3xflag*]) is a gift from Feldman lab. To generate ***shrEx483*,** young adult worms were microinjected with 5 ng/µL of *pser2prom3::gfp::klp-7b*, 10 ng/µL of *pser2prom3::mScarlet*, and 50 ng/µL of *pttx-3::rfp*. The construct of *pser2prom3::gfp::klp-7b* was generated using the InFusion® cloning.

### Sample preparation

Neuronal polarity was assayed in L4 staged worms for both control and mutant conditions. The worms were harvested after a definite period and are mounted on 7.5% agarose pads using a cocktail of 0.1% Tricaine and 10mM Tetramisole. Similar preparation was done for other reporters like PTRN-1::tagRFP and UNC-44::GFP.

For imaging of microtubule-associated End Binding Protein (EBP) comets, the worms were mounted on 10% agarose pads in a mounting medium of 0.1µm polystyrene beads. A maximum of 5-7 worms were mounted simultaneously for imaging. For FRAP assays, the mounting was done in a mixture of 0.1% Tricaine and 10 mM Tetramisole as described before.

Worms were grown on Colchicine (1mM) containing NGM plates since hatching for the pharmacological perturbation of the microtubules.

### Microscopy

Worms were imaged on the Nikon A1R confocal using 60X/1.4NA oil objective at a voxel scale of 0.414 µm x 0.414 µm x 1.0 µm. A large area scanning was enabled to image the whole worm. To image multiple reporters sequential channel imaging was used. For some ubiquitously expressed reporters like UNC-44::GFP, the imaging was carried out at a voxel scale of 0.09 µm x 0.09 µm x 0.2 µm using 60X/1.4NA oil objective.

Live imaging of EBP comets was carried out on the Zeiss Observer.Z1 inverted microscope equipped with Yokogawa CSU-X1 spinning disk unit and Photometrics Evolve camera. Imaging was carried out on a 100X/1.4NA oil objective at a temporal resolution of 1 frame per second for a duration of 2 min.

FRAP assays were carried out on the Nikon A1R confocal using 60X/1.4NA oil objective at a voxel scale of 0.09 µm x 0.09 µm x 0.2 µm. A circular region of 5 µm diameter was chosen for the photobleaching and the entire acquisition was carried out at a frame rate of 1 frame per second. Before photobleaching 5 frames were collected, photobleaching was carried out for 5 seconds and post photobleaching acquisition was carried out for a duration of 2 minutes.

### Analysis

Neuronal polarity has been assayed by the percentage of dendrites with mCherry::RAB-3, density of mCherry::RAB-3 punctae, and population of worms with dendritic SAD-1::GFP in the primary dendrites. The percentage of dendrites and population distribution has been assessed manually by locating punctae over a particular threshold intensity. This threshold has been set by the intensity of the mCherry::RAB-3 punctae present in the axon (ventral nerve cord) in the wildtype neurons. The density of mCherry::RAB-3 punctae has been assessed by the line profile of intensities in the primary dendrites and peak finding tool using the BAR plugin in ImageJ. PTRN-1::tagRFP punctae were also analyzed for their density in the primary dendrites using the line profile and peak finding tool using the BAR plugin in ImageJ.

Time lapse acquisitions of EBP-2::GFP comets were analyzed in ImageJ by extracting kymographs from different neurites of PVD neurons. The traces in the kymographs were assessed using the line tool to obtain the angle, length, and duration. Slopes of the traces obtained from the angles was used for the orientation of the comets. Length of the traces was divided by their duration to obtain the rates of polymerization.

The images and videos were quantified using ImageJ®. All the images were background corrected and normalized to the maximum. Intensities obtained from the FRAP region were normalized to pre-bleach and post-bleach intensities and plotted against time. The FRAP curves were fitted to the following single exponential equation using the Levenberg-Marquardt iteration algorithm in Origin®:

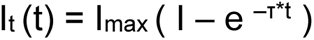

The maximum recovery amplitude (I_max_) and the half-time (t_1/2_ = ln0.5/-τ) parameters were obtained from the fit.

Intensities in a given neurite were quantified from a line ROI spanning across multiple optical sections and normalized with a constitutive marker.

Quants obtained from ImageJ® were fed into Excel and Origin® to get statistically relevant data. Parametric data were compared using ANOVA and Bonferroni test. Nonparametric datasets were compared using Fisher’s test.

## Acknowledgments

We thank Dr. Jessica Feldman for the CRISPR allele of EBP-2::GFP and Caenorhabditis Genetics Center (CGC) for mutant and transgenic strains. CGC is supported by the NIH Office of Research Infrastructure Programs (P40 OD010440). This work is supported by the NBRC core fund from the Department of Biotechnology, and DBT/Wellcome Trust India Alliance (Grant # IA/E/13/1/504331 to S.D., Grant # IA/I/13/1/500874 to A.G.-R.).

## Author Contributions

S.D., N.K. and A.G-R designed experiments. S.D., and N.K. performed experiments and analyzed data. S.D. and A.G-R wrote and edited the manuscript. J.F. shared the EBP-2::GFP reagent and provided suggestions for microtubule assays.

