## Supplemental Data for "KLP-7/Kinesin-13 orchestrates axon-dendrite checkpoints for polarized trafficking in neurons"

**List of Figures:**

**Figure S1 (linked to Figure 1): Spatial distribution of mCherry::RAB-3 in the dendrites depends on KLP-7.**

**Figure S2 (linked to Figure 2): Effect of the Colchicine treatment on the axon-dendrite compartmentalization.**

**Figure S3 (linked to Figure 3): KLP-7 regulates the microtubule organization in PVD neurites.**

**Figure S4 (linked to Figure 4): Effect of *klp-7(0)* on the levels and expression of UNC-44::GFP in the PVD neurons*.***

**Figure S5 (linked to Figure 5): KLP-7 is more dynamic in the dendrites than in the axon of the PVD neurons.**

**List of Tables:**

**Table S1: List of strains and constructs.**

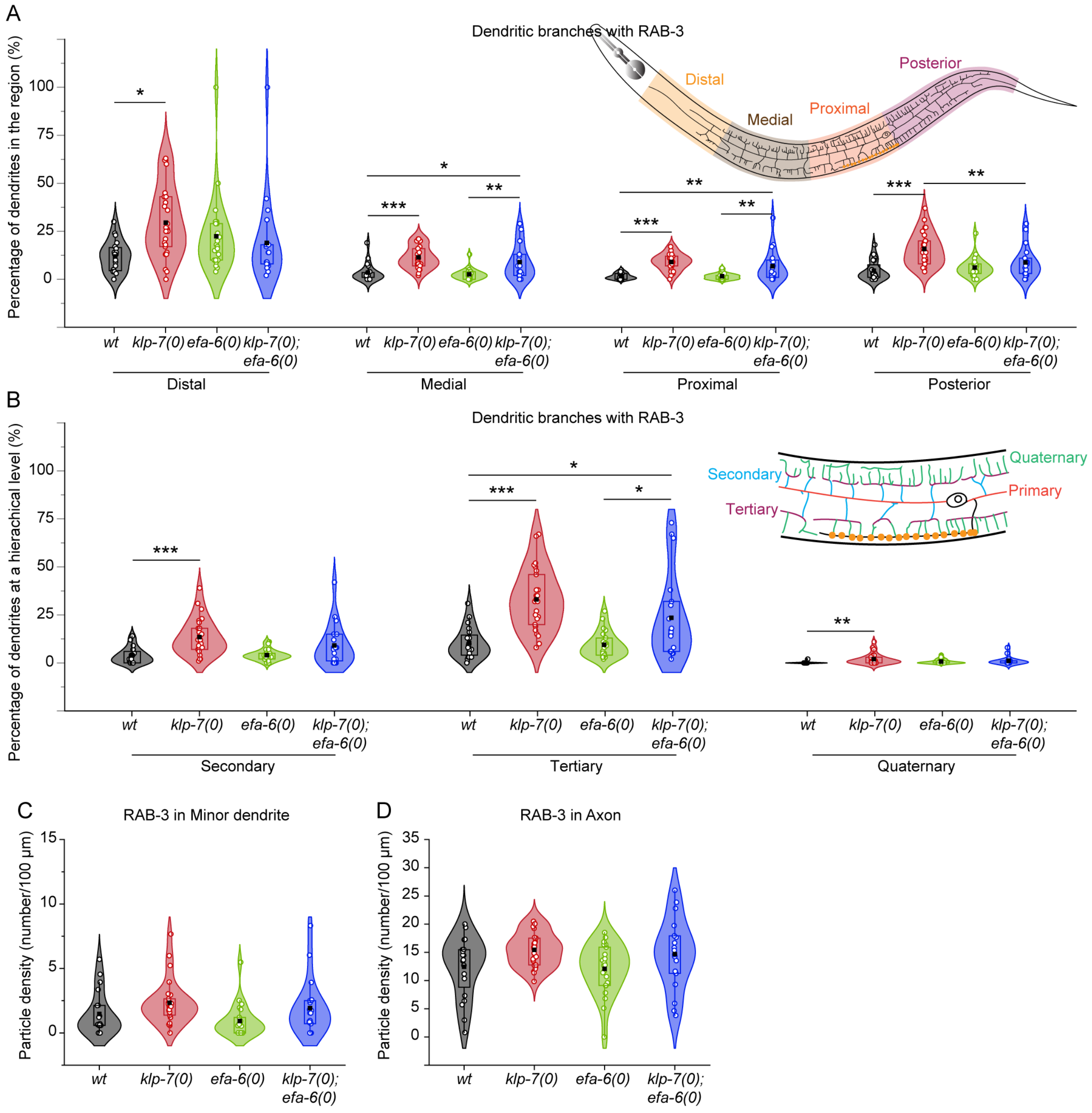


**Figure S1 (linked to Figure 1): Spatial distribution of mCherry::RAB-3 in the dendrites depends on KLP-7.**

A. Schematic shows the demarcation of the distal, medial, and proximal regions which are one third of the anterior dendrite and the posterior dendrite. Percentage of dendrites in the distal, medial, proximal, and posterior regions of the PVD neuron quantified for the presence of mCherry::RAB-3 in wildtype (wt), *klp-7(0)*, *efa-6(0)*, and *klp-7(0);efa-6(0)* double mutant. Region wise comparison of means was done using ANOVA and Bonferroni Test. $p < 0.05$*, 0.01**, 0.001***.

B. Schematic shows the branch order in the PVD neuron quantified for the mCherry::RAB-3 accumulation in each hierarchical level in wildtype (wt), *klp-7(0)*, *efa-6(0)*, and *klp-7(0);efa-6(0)* double mutant. Branch order wise comparison of means was done using ANOVA and Bonferroni Test. $p < 0.05$*, 0.01**, 0.001***.

C-D. Particle density of mCherry::RAB-3 quantified for the posterior dendrite (minor), and the axon of PVD neurons of wildtype (wt), *klp-7(0)*, *efa-6(0)*, and *klp-7(0);efa-6(0)* double mutant. Comparison of means was done using ANOVA and Bonferroni Test and $p > 0.05$.

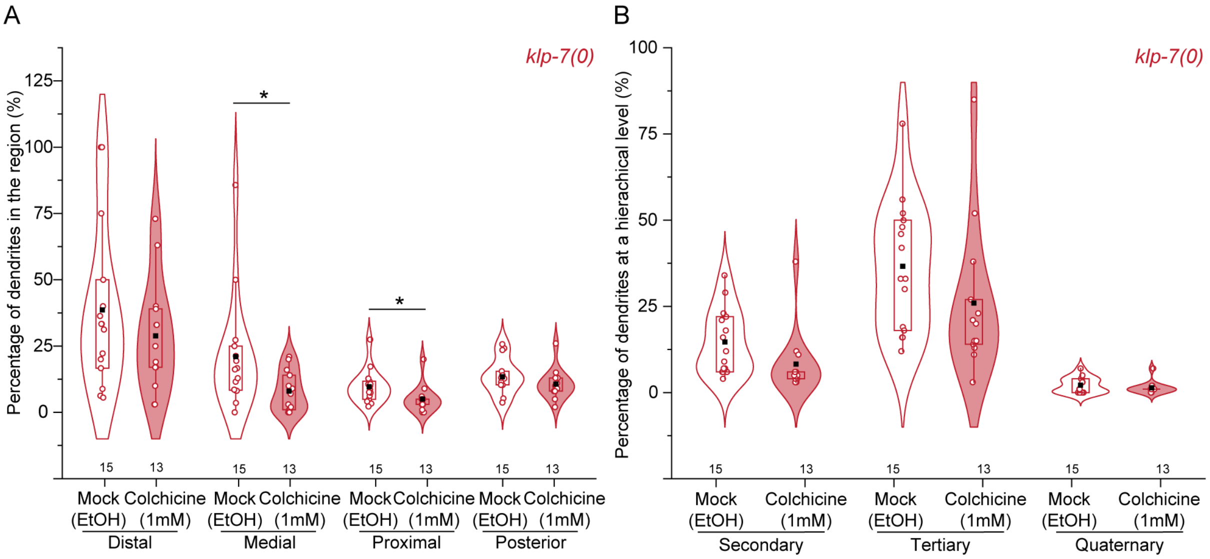


**Figure S2 (linked to Figure 2): Effect of the Colchicine treatment on the axon-dendrite compartmentalization.**

A-B. Percentage of dendrites in different regions viz. distal, medial, proximal, and posterior (A) and hierarchical levels including secondary, tertiary, and quaternary (B) of the PVD neuron in *klp-7(0)* mutant upon Colchicine treatment (F) with their respective mock controls. Comparison of means was done using ANOVA and Bonferroni Test and $p > 0.05$. Number of animals assessed (n) is mentioned along the X-axis.

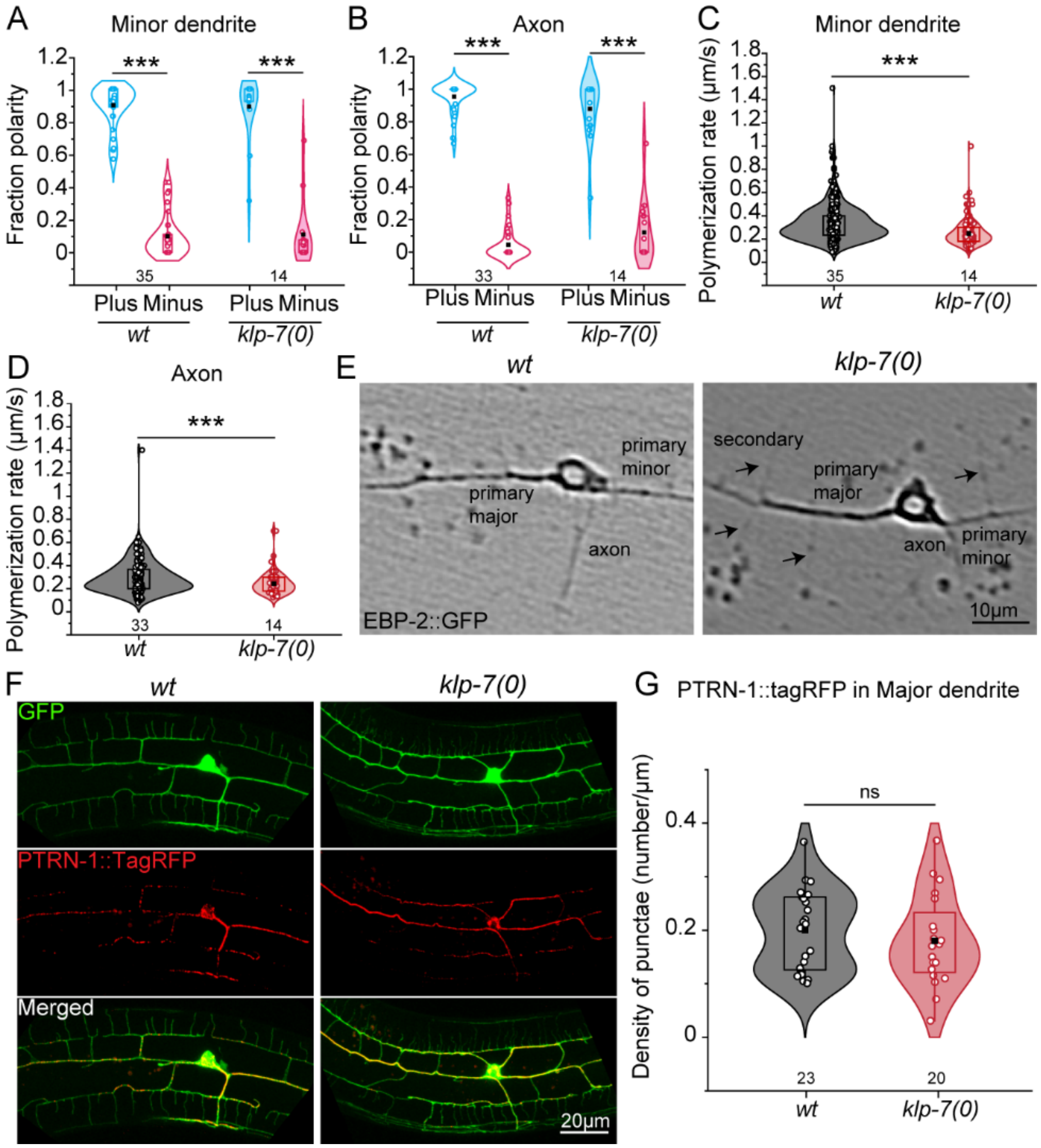


**Figure S3 (linked to Figure 3): KLP-7 regulates the microtubule organization in PVD neurites.**

A-B. Relative orientation of the microtubules is represented as a fraction polarity of EBP-2::GFP comets in the plus-end-out (cyan) and minus-end-out directions (magenta) in the minor dendrite (A) and axon (B) of the wildtype (wt) and *klp-7(0)* PVD neurons. Comparison of means was done using ANOVA and Bonferroni Test. $p < 0.001$***. Number of animals assessed (n) is mentioned along the X-axis.

C-D. Rates of polymerization were quantified as the covered distance divided by the duration of the EBP-2::GFP comets in the minor dendrite (C) and axon (D) of wildtype (wt) and *klp-7(0)*. Comparison of means was done using ANOVA and Bonferroni Test. $p < 0.001$***. Number of animals assessed (n) is mentioned along the X-axis.

E. Distribution of EBP-2::GFP expressed under PVD promoter in various neurites of the PVD neuron of wildtype (wt) and *klp-7(0)*.

F-G. Representative images and quantification of PTRN-1::TagRFP punctae expressed in the PVD neuron in the wildtype (wt) and *klp-7(0)*. Comparison of means was done using ANOVA and Bonferroni Test. ns, (not significant). Number of animals assessed (n) is mentioned along the X-axis.

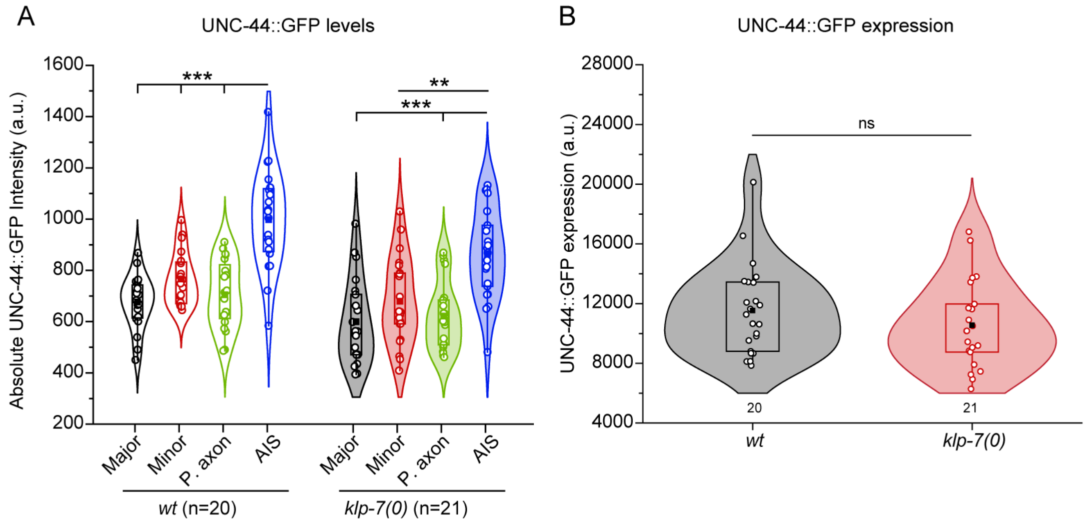


**Figure S4 (linked to Figure 4): Effect of *klp-7(0)* on the levels and expression of UNC-44::GFP in the PVD neurons*.***

A. Intensities of UNC-44::GFP compared between major, minor, proximal axon (P. axon) and Axon Initial Segment (AIS) in the wildtype (wt) and *klp-7(0)*. Comparison of means was done using ANOVA and Bonferroni Test. p<0.01**, 0.001***. Number of animals assessed (n) is mentioned along the X-axis.

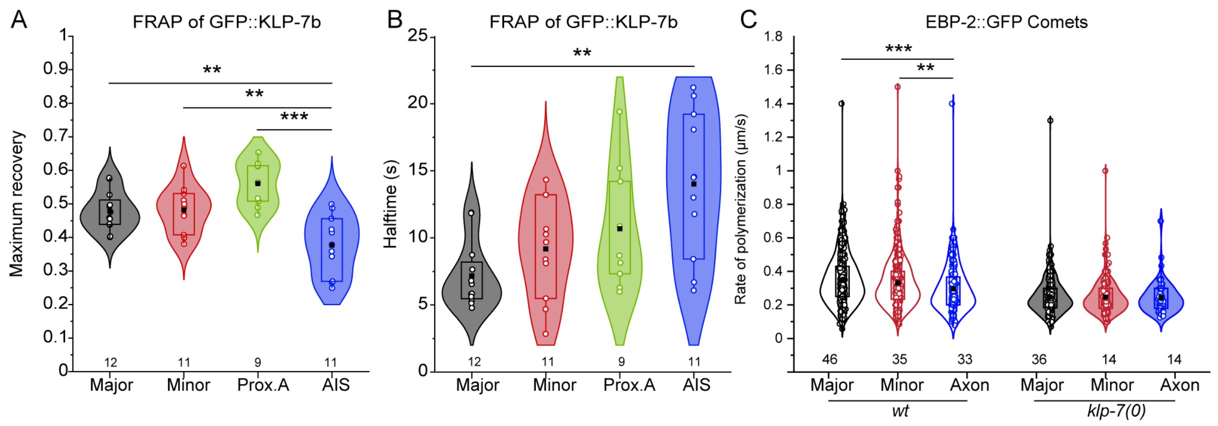


**Figure S5 (linked to Figure 5): KLP-7 is more dynamic in the dendrites than in the axon of the PVD neurons.**

A-B. Maximum recovery ratio (A) and half times of the FRAP recovery profiles of GFP::KLP-7b in various compartments of the PVD neurons. Statistical comparison was done using ANOVA and Bonferroni Test. p < 0.01**, 0.001***. Number of animals assessed (n) is mentioned along the X-axis.

C. Rates of polymerization obtained from the EBP-2::GFP comets in the various compartments of the PVD neurons in the wildtype (wt) and *klp-7(0)* animals. Statistical comparison between the compartments was done using ANOVA and Bonferroni Test. p < 0.01**, 0.001***. Number of animals assessed (n) is mentioned along the X-axis.

**Table S1: List of strains and constructs.**

| Reagent type (species) or resource | Designation | Source or reference | Identifiers | Additional information |
|---|---|---|---|---|
| Strain, *C.elegans* | ***kyIs445*** *[pdes-2::mCherry::RAB-3; pdes-2::SAD-1::GFP; podr-1::DsRed]* *;* ***wdIs52****[pF49H12.4::GFP + unc-119(+)]* | (Puri *et al.*, 2021) | NBR374 | ***kyIs445*** *[pdes-2::mCherry::RAB-3 (0.5 ng/µL); pdes-2::SAD-1::GFP (2 ng/µL); podr-1::DsRed (30 ng/µL)],* a gift from Bargmann lab, integrated extrachromosomal array ***wdIs52****[pF49H12.4::GFP + unc-119(+)]*, integrated on X-chromosome for PVD specific expression of GFP |
| Strain, *C.elegans* | ***klp-7(tm2143)*** *;* ***kyIs445****[pdes-2::mCherry::RAB-3; pdes-2::SAD-1::GFP; podr-1::DsRed]* *;* | (Puri *et al.*, 2021) | NBR375 | ***klp-7(tm2143)***, gene knockout with 875bp deletion ***kyIs445*** *[pdes-2::mCherry::RAB-3 (0.5 ng/µL); pdes-2::SAD-1::GFP (2 ng/µL); podr-1::DsRed (30* |

| | | | | |
|---|---|---|---|---|
| | ***wdIs52***[*pF49H12.4::GFP + unc-119(+)*] | | | *ng/µL)],* a gift from Bargmann lab, integrated extrachromosomal array ***wdIs52***[*pF49H12.4::GFP + unc-119(+)*], integrated on X-chromosome for PVD specific expression of GFP |
| Strain, *C.elegans* | ***efa-6(ok3533)*** ; ***kyIs445***[*pdes-2::mCherry::RAB-3; pdes-2::SAD-1::GFP; podr-1::DsRed*] ; ***wdIs52***[*pF49H12.4::GFP + unc-119(+)*] | Crossed in this study | NBR1007 | ***efa-6(ok3533)***, gene knockout mutant with an estimated deletion of 700bp (Barstead *et al.*, 2012) ***kyIs445*** *[pdes-2::mCherry::RAB-3 (0.5 ng/µL); pdes-2::SAD-1::GFP (2 ng/µL); podr-1::DsRed (30 ng/µL)],* a gift from Bargmann lab, integrated extrachromosomal array ***wdIs52***[*pF49H12.4::GFP + unc-119(+)*], integrated on X-chromosome for PVD specific expression of GFP |
| Strain, *C.elegans* | ***klp-7(tm2143)*** ; ***efa-6(ok3533)*** ; ***kyIs445***[*pdes-* | Crossed in this study | NBR1034 | ***klp-7(tm2143)***, gene knockout with 875bp deletion |

| | | | | |
|---|---|---|---|---|
| | *2::mCherry::RAB-3; pdes-2::SAD-1::GFP; podr-1::DsRed] ;* ***wdIs52****[pF49H12.4::GFP + unc-119(+)]* | | | ***efa-6(ok3533)***, gene knockout mutant with an estimated deletion of 700bp (Barstead *et al.*, 2012) ***kyIs445*** *[pdes-2::mCherry::RAB-3 (0.5 ng/µL); pdes-2::SAD-1::GFP (2 ng/µL); podr-1::DsRed (30 ng/µL)],* a gift from Bargmann lab, integrated extrachromosomal array |
| Strain, *C.elegans* | *ebp-2*(***wow47***[*ebp-2::gfp::3xflag*]) *;* ***shrEx472****[pser2prom3::mscarlet; pttx3::RFP]* | Crossed in this study | NBR225 | *ebp-2*(***wow47***[*ebp-2::gfp::3xflag*]) gift from Feldman lab, GFP and 3xFlag tags inserted into the C-terminus of the endogenous *ebp-2* locus ***shrEx472****[pser2prom3::mscarlet (10 ng/µL);pttx3::RFP (50 ng/ µL]* generated in Ghosh-Roy lab, Extrachromosomal array with transmission~90% |
| Strain, *C.elegans* | *ebp-2*(***wow47***[*ebp-* | Crossed in this study | NBR228 | *ebp-2*(***wow47***[*ebp-2::gfp::3xflag*]) gift from Feldman lab, GFP and 3xFlag |

| | | | | |
|---|---|---|---|---|
| | *2::gfp::3xflag*]) *;* ***klp-7(tm2143)*** *;* ***shrEx472****[pser2prom3::mscarlet; pttx3::RFP]* | | | tags inserted into the C-terminus of the endogenous *ebp-2* locus ***klp-7(tm2143)***, gene knockout with 875bp deletion ***shrEx472****[pser2prom3::mscarlet (10 ng/µL);pttx3::RFP (50 ng/ µL]* generated in Ghosh-Roy lab, Extrachromosomal array with transmission~90% |
| Strain, *C.elegans* | *unc-44(****hrt5****[GFPKI])* *;* ***shrEx472****[pser2prom3::mscarlet; pttx3::RFP]* | Crossed in this study | NBR226 | STR282:*unc-44(****hrt5****[GFPKI])* obtained from Caenorhabditis Genetics Centre (CGC) (He *et al.*, 2020) ***shrEx472****[pser2prom3::mscarlet (10 ng/µL);pttx3::RFP (50 ng/ µL]* generated in Ghosh-Roy lab, Extrachromosomal array with transmission~90% |
| Strain, *C.elegans* | *unc-44(****hrt5****[GFPKI])* *;* ***klp-7(tm2143)*** *;* ***shrEx472****[pser2pr* | Crossed in this study | NBR418 | STR282:*unc-44(****hrt5****[GFPKI])* obtained from Caenorhabditis Genetics Centre (CGC) (He *et al.*, 2020) |

| | | | | |
|---|---|---|---|---|
| | *om3::mscarlet; pttx3::RFP]* | | | ***klp-7(tm2143)***, gene knockout with 875bp deletion ***shrEx472****[pser2prom3::mscarlet (10 ng/µL);pttx3::RFP (50 ng/ µL]* generated in Ghosh-Roy lab, Extrachromosomal array with transmission~90% |
| Strain, *C.elegans* | ***shrEx483****[pser2prom3::gfp::klp-7b;pser2prom3::mScarlet;pttx-3::rfp]* | Made in this study | NBR383 | ***shrEx483****[pser2prom3::gfp::klp-7b (5 ng/µL);pser2prom3::mScarlet (10 ng/µL);pttx-3::rfp (50 ng/µL)]* Extrachromosomal array with transmission~86.4% |
| Strain, *C.elegans* | ***shrEx307****[pser2::ebp-2::gfp; pttx3::RFP]* | Made in this study | NBR716 | ***shrEx307****[pser2::ebp-2::gfp (5ng/µL); pttx3::RFP(50 ng/µL)]* Extrachromosomal array with transmission~75% |
| Strain, *C.elegans* | ***klp-7(tm2143)*** *;* ***shrEx307****[pser2::ebp-2::gfp; pttx3::RFP]* | Crossed in this study | NBR714 | ***klp-7(tm2143)***, gene knockout with 875bp deletion ***shrEx307****[pser2::ebp-2::gfp (5ng/µL); pttx3::RFP(50 ng/µL)]* Extrachromosomal array with transmission~75% |

| | | | | |
|---|---|---|---|---|
| DNA construct | *pser2prom3::**gfp::klp-7b*** | Made in this study | pNBR47 | InFusion® cloning |
| Oligos | *Forward primer for amplification of vector backbone containing **pser2prom3*** | | AGR644 | CAGCTTTCTTGTACAAAGTGGTGA |
| Oligos | *Reverse primer for amplification of vector backbone containing **pser2prom3*** | | AGR645 | AAGGGCGAATTCGGAGCCT |
| Oligos | *Forward primer for amplification of insert containing **gfp::klp-7b*** | | AGR646 | TCCGAATTCGCCCTTATGAGTAAAGGAGAAGAACTTTTCA |
| Oligos | *Reverse primer for amplification of insert containing **gfp::klp-7b*** | | AGR647 | TGTACAAGAAAGCTGTCAGACGTTTTCCACGGCGA |
| Oligos | *Forward primer for the genotyping of klp-7(tm2143)* | | AGR179 | CGGCAGTACTCGCCATTTCATAAC |

| | | | | |
|---|---|---|---|---|
| Oligos | *Internal forward primer for the genotyping of klp-7(tm2143)* | | AGR181 | TCTCACGCTGAACCCATCACTC |
| Oligos | *Reverse primer for the genotyping of klp-7(tm2143)* | | AGR180 | CATCGACGCATTCTGCTTCTTTCC |
| Oligos | *Forward primer for the genotyping of efa-6(ok3533)* | | AGR836 | GATCATTGGACGCAAAGGATATGTCA |
| Oligos | *Internal forward primer for the genotyping of efa-6(ok3533)* | | AGR848 | AGCAATAGTTCAGCATCAGCCAG |
| Oligos | *Reverse primer for the genotyping of efa-6(ok3533)* | | AGR849 | ACTGAACCATTTGCCGTCGAG |
